# Is genetic liability to ADHD and ASD causally linked to educational attainment?

**DOI:** 10.1101/2020.02.11.944041

**Authors:** Christina Dardani, Beate Leppert, Lucy Riglin, Dheeraj Rai, Laura D Howe, George Davey Smith, Kate Tilling, Anita Thapar, Neil M. Davies, Emma Anderson, Evie Stergiakouli

**Affiliations:** Centre of Academic Mental Health, Bristol Medical School, University of Bristol, UK; Medical Research Council Integrative Epidemiology Unit, Bristol Medical School, University of Bristol, UK; Population Health Sciences, Bristol Medical School, University of Bristol, UK; Division of Psychological Medicine and Clinical Neurosciences, MRC Centre for Neuropsychiatric Genetics and Genomics, Cardiff University, UK

## Abstract

**Background:** Individuals with Attention Deficit Hyperactivity Disorder (ADHD) or Autism Spectrum Disorder (ASD) are at risk of poor educational outcomes. Parental educational attainment has also been associated with risk of ADHD/ASD in the offspring. Despite evidence that ADHD and ASD show genetic links to educational attainment, less is known on the causal nature of the associations and the possible role of IQ.

**Methods:** We assessed the total causal effects of genetic liability to ADHD/ASD on educational attainment using two-sample Mendelian randomization (MR). We assessed the possible contribution of IQ to the identified causal effects by estimating the “direct” effects of ADHD/ASD on educational attainment, independent of IQ, using Multivariable MR (MVMR). Reverse direction analyses were performed. The latest GWAS meta-analyses of ADHD, ASD, educational attainment and IQ were used. Causal effect estimates were generated using inverse variance weighted models (IVW). Sensitivity analyses were performed to assess the robustness of the estimates and the presence of pleiotropy.

**Results:** Genetic liability to ADHD had a total (_MR_IVW:-3.3 months per doubling of liability to ADHD; _95%_CI: -4.8 to -1.9; pval= 5*10^−6^) and direct negative causal effect on educational attainment (_MVMR_IVW:-1.6 months per doubling of liability to ADHD; _95%_CI: -2.5 to -0.6; pval= 4*10^−4^). There was little evidence of a total causal effect of genetic liability to ASD on educational attainment (_MR_IVW: 4 days, per doubling of liability to ASD; _95%_CI: -4.9 months to 5.6 months; pval= 0.9) but some evidence of a direct effect not via IQ (_MVMR_IVW:29 days per doubling the genetic liability to ASD; _95%_CI: 2 to 48; pval= 0.03). Reverse direction analyses suggested that genetic liability to higher educational attainment was associated with lower risk of ADHD (_MR_IVW_OR_: 0.3 per standard deviation (SD) increase; _95%_CI: 0.26 to 0.36; pval= 6*10^−51^), even after IQ was entered in the models (_MVMR_IVW_OR:_ 0.33 per SD increase; _95%_CI: 0.26 to 0.43; pval= 6*10^−17^). On the contrary, there was evidence consistent with a positive causal effect of genetic liability to higher educational attainment on risk of ASD (_MR_IVW_OR_: 1.51 per SD increase; _95%_CI: 1.29 to 1.77; pval= 4*10^−7^), which was found to be largely explained by IQ (_MVMR_IVW_OR_ per SD increase: 1.24; _95%_CI: 0.96 to 1.60; pval= 0.09).

**Conclusions:** Our findings suggest that despite the genetic and phenotypic overlap between ADHD and ASD, they present highly differentiated causal associations with educational attainment. This highlights the necessity for specialized educational interventions for children with ADHD and ASD. Further research is needed in order to decipher whether the identified causal effects reflect parentally transmitted effects, diagnostic masking, or selection bias.

## INTRODUCTION

Attention Deficit Hyperactivity Disorder (ADHD) and Autism Spectrum Disorder (ASD) are neurodevelopmental conditions, that typically first manifest early in childhood and often persist into adulthood^1,2^. Both conditions are associated with one of the strongest predictors of adult life outcomes and life satisfaction; educational attainment^3,4^.

Observational research evidence suggests that children with ADHD show lower academic performance compared to their typically developing peers^5–7^ and the condition has been also associated with increased risk of high school dropout^8^. In the case of ASD, rates of transition to post-secondary education are much lower than in the general population, and only a small proportion of individuals with ASD who move on to higher education will eventually graduate^9,10^. Several factors have been found to predict educational attainment in children with ADHD or ASD, with one of the strongest being IQ^11,12^. The pattern of association of ADHD and ASD with educational outcomes is further complicated by parental educational attainment. Cohort and registry-based studies suggest that parental educational attainment and socioeconomic position are associated with risk of ADHD and ASD in the offspring, in opposite directions (higher educational attainment associated with increased risk of ASD; lower educational attainment associated with increased risk of ASD)^13,14^. Observational evidence has been recently corroborated by studies utilizing whole genome approaches (LD score regression^15^, MTAG^16^), as well as aggregates of common risk variants (i.e. polygenic risk scores (PRS), suggesting strong negative genetic correlations of educational attainment with ADHD and positive with ASD^17–19^.

Despite their genetic and phenotypic overlap, the causal nature of these associations has not been investigated. Observational evidence can be hampered by measured or unmeasured confounding^20^ and whole genome approaches and PRS do not account for the potential influence of pleiotropic genetic variants^21^. This is particularly important when analysing traits that are highly phenotypically and genetically correlated (for example, education and IQ). A useful method for overcoming these limitations is Mendelian randomization (MR). MR is a form of instrumental variables analysis, utilizing common genetic variants as proxies for environmental exposures and allowing the assessment of causal relations among the exposures with the outcome of interest^22^. The method, under certain assumptions, has the potential to unpick the direction of the causal effect and investigate the presence of pleiotropy^23^. Several possible causal pathways linking ADHD, ASD, IQ and educational attainment could be proposed-some of them visualised in Figure 1. Similar possibilities could exist for the causal pathways linking educational attainment to ADHD/ASD. Deciphering the possible contribution of IQ in the causal pathways linking genetic liability to ADHD, ASD and educational attainment is an important step towards designing targeted interventions and offering adequate educational and psycho-social support in order to improve the quality of life and adult life outcomes of the affected individuals.

**Figure 1.**
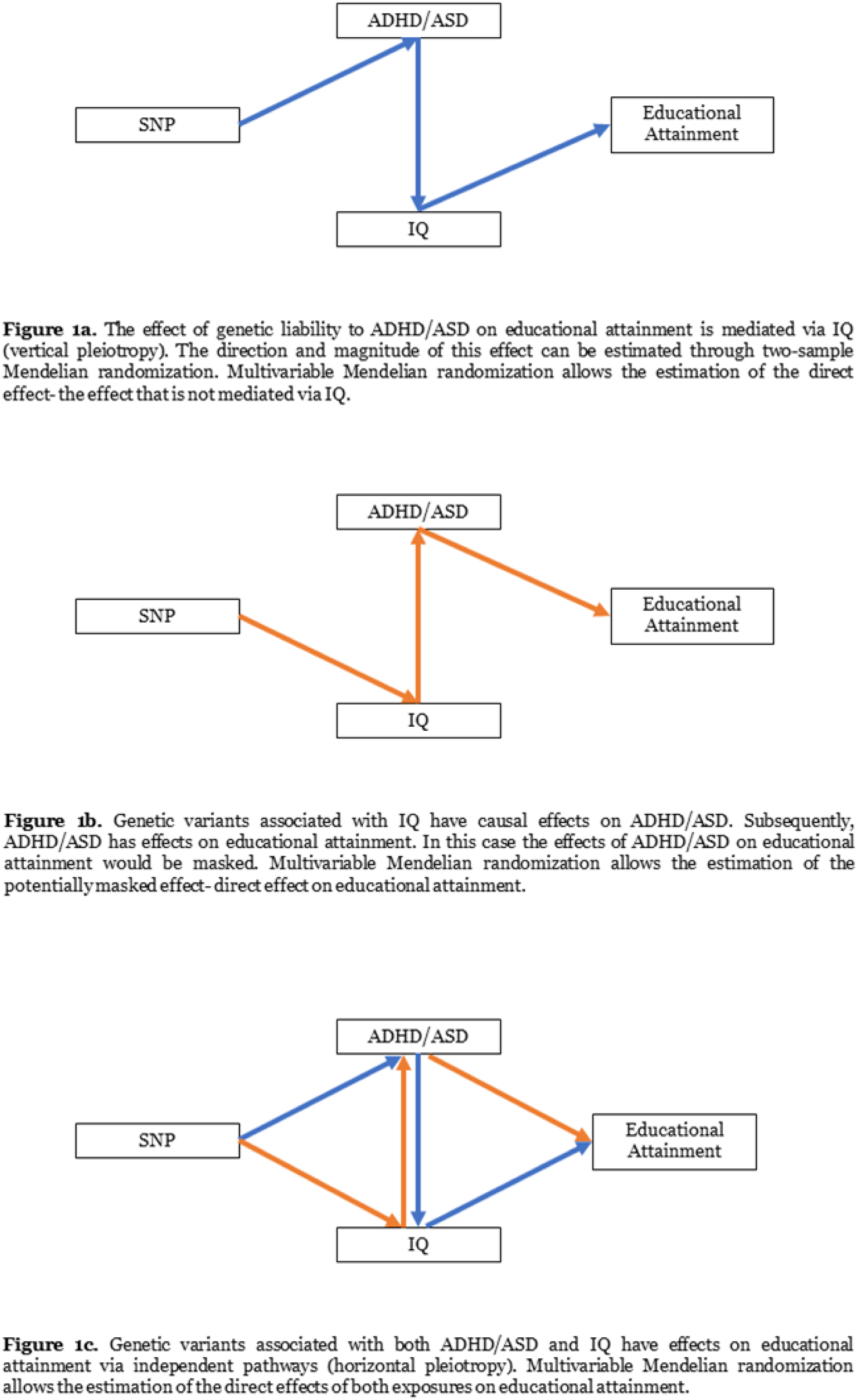
Possible causal pathways linking ADHD/ASD, IQ and educational attainment. The figures are not directed acyclic graphs, do not cover all the causal pathways that might exist between the phenotypes and do not imply that ADHD and ASD share the same causal links to educational attainment.

In the present study, we aimed to assess whether:

i. genetic liability to ADHD and ASD has a causal impact on educational attainment
ii. genetic liability to higher educational attainment (reflecting parental effects) has a causal impact on risk of ADHD and ASD,

in a two-sample MR framework. An extension of MR, Multivariable MR (MVMR), was used to assess the possible role of genetic liability to ADHD and ASD that is not mediated via IQ^24^.

## METHODS

### Univariable two-sample Mendelian randomization

MR allows the estimation of causal associations between an exposure and an outcome by utilizing common genetic variants as instruments for the exposure of interest. The robustness of the method relies on strict assumptions that the genetic instruments should satisfy: (i) there must be a robust association between the implicated genetic variants and the exposure, (ii) the variants should operate on the outcome entirely via the exposure, (iii) the variants should not be associated with any confounders of the associations between the exposure and the outcome^25^. In this context, we applied two-sample MR in which the effects of the genetic instruments on the exposure and on the outcome are extracted from separate GWASs that have been conducted in independent samples from the same underlying population^22^.

### Genetic instruments

We used the latest publicly available GWAS summary statistics on ADHD^17^, ASD^18^, IQ^26^ and educational attainment^27^. Detailed information on the GWASs used can be found in the original publications.

In each GWAS dataset, we extracted all the variants with a p-value <= 5*10^−8^. The identified variants were clumped using an r^2^< 0.01, within a 10,000 kb window, based on the 1000Genomes European phase 3 reference panel. This resulted in 11 SNPs in ADHD, 2 SNPs in ASD, 481 SNPs in educational attainment and 212 SNPs in IQ.

In order to increase the power of the ASD analyses, we relaxed the p-value threshold to 5*10-^7^. After clumping, we identified 10 independent (r^2^< 0.01) SNPs. A similar threshold (p<= 5*10^−6^) for instrument definition has been used in previous studies^28,29^. Details on the effect sizes, standard errors and p-values of the instruments can be found in Supplementary table S1.

For each analysis, instruments were extracted from the outcome GWAS. LD link was used to identify LD proxies, when SNPs were not present in the outcome GWAS (r^2^>0.9).

Finally, the alleles of the outcome variants were harmonised on the exposure so that the effect estimates of both exposure and outcome variants were expressed per increasing allele. As the effect allele frequencies for the ADHD and ASD GWASs were not provided, when the harmonisation of the exposure-outcome alleles was not possible, variants were excluded from the analyses as being palindromic. Detailed information on the harmonised datasets used in the present MR analyses can be found in Supplementary table S2. The full process followed, and the final number of instruments used for each analysis is visualised in Supplementary Figure S1.

Two-sample MR analyses were performed using the TwoSampleMR R package^30^.

### Inverse Variance Weighted MR

The primary MR method used in this study was the Inverse Variance Weighted (IVW) regression. It is a weighted generalised linear regression of the SNP-outcome coefficients on the SNP-exposure coefficients with a constrained intercept term, giving an overall causal effect estimate of the exposure on the outcome^31^.

### Instrument Strength

We assessed the strength of the instruments by estimating the mean F-statistic as a cumulative estimate of weak instrument bias influencing the IVW. As a rule of thumb, if the mean F > 10, then the IVW is unlikely to suffer from weak instrument bias^32^.

### Sensitivity Analyses

To test for the presence of horizontal pleiotropy (i.e. whereby a genetic variant has independent effects on multiple phenotypes) we used MR-Egger regression^31,33^. MR Egger (in contrast to IVW) allows the intercept term to be unconstrained. The intercept parameter indicates the overall unbalanced horizontal pleiotropic effect of the SNPs on the outcome (i.e. a direct effect of each SNP on the outcome, independent of the exposure, which would violate MR assumptions), while the slope offers a causal effect estimate accounting for directional pleiotropy. MR-Egger, as all MR analyses, assumes the gene-exposure association estimates are measured without error (i.e. the no measurement error [NOME] assumption)^31^. We assessed the NOME assumption using an adaptation of the I^2^ statistic^34^ within the two-sample MR context, which is referred to as I^2^_GX_^33^.I^2^_GX_ provides an estimate of the degree of regression dilution in the MR-Egger causal estimate, due to uncertainty in the SNP-exposure estimates. We then used simulation extrapolation (SIMEX) to adjust the MR-Egger estimate for this dilution, as described previously^33^. We conducted weighted median MR which provides an estimate of causal effect even when only 50% of the genetic variants included in the analysis are valid instruments for the exposure^35^.

Finally, we applied Steiger filtering to assess whether each SNP explained more variation in the exposure that is instrumenting, than in the outcome^36^. This way, we scrutinised whether the direction of the estimated causal effect was correct and not influenced by the large sample size of the educational attainment GWAS or the strong genetic associations between ADHD, ASD and educational attainment^17,18^.

### Multivariable Mendelian randomization

Where multiple exposures are suspected to have causal effects on an outcome, and the exposures are genetically and phenotypically correlated, univariable MR can yield biased causal effect estimates^24^. Multivariable MR (MVMR) is an extension of MR, in which multiple exposures are entered within the same model, and their direct effects on the outcome can be estimated^24^. We used MVMR to estimate the direct effects of ADHD, ASD and IQ on educational attainment and the direct effects of genetic liability to higher educational attainment and IQ on risk of ADHD and ASD.

For each MVMR analysis, 212 genome-wide significant and independent (r^2^<0.01, 10.000kb window) instruments for IQ were entered in the models. The full list of primary exposure instruments and IQ instruments was clumped (r^2^=0.01), to ensure the absence of LD among the included SNPs, and then harmonised. We performed an inverse variance weighted (IVW) regression of the SNP-outcome coefficients on the SNP-exposure coefficients, entering the two exposures in the regression model simultaneously. Details on the effect sizes, standard errors and p-values of the IQ instruments used, can be found in Supplementary table S1. We estimated the heterogeneity of the causal effect estimates of the instruments included, using a modified version of the Q-statistic in the context of MVMR. Evidence of heterogeneity indicates the possibility of biased causal effect estimates^24^. Additionally, as a sensitivity analysis, in cases that Steiger filtering suggested that SNPs explained more variation in the outcome than in the exposure, we repeated MVMR analyses by removing these SNPs.

Details on the MVMR method and analytic process have been described elsewhere^24^. MVMR analyses were performed using R, version 3.5 and Q statistics were estimated using the MVMR package (https://github.com/WSpiller/MVMR).

### Interpretation of the causal effect estimates

In the analyses investigating the causal effects of genetic liability to ADHD, ASD on educational attainment, the estimated MR effects and 95% confidence intervals were expressed as one standard deviation (SD) increase in educational attainment per log-odd increase in genetic liability to ADHD, ASD. In order to assist the interpretation of the findings, we converted the estimates to months/days (where appropriate) of education per doubling the genetic liability to ADHD and ASD. Specifically, we multiplied the estimates by the standard deviation of educational attainment (years of education SD = 3.8099)^27^ and then multiplied by ln(2) to estimate the effect of a doubling of liability to ADHD, ASD.

In the analyses investigating the causal effects of genetic liability to higher educational attainment on risk of ADHD and ASD, MR estimates and 95% confidence intervals are expressed per one SD increase in educational attainment on the odds of developing ADHD, ASD.

## RESULTS

### Total causal effect of genetic liability to ADHD on educational attainment

The mean F-statistic of the ADHD instruments was ≈ 35. The IVW causal effect estimate suggested that a doubling in the genetic liability to ADHD decreases years of education by around 3 months (_MR_IVW:-3.3 months per doubling the liability to ADHD; _95%_CI: -4.8 to - 1.9; pval= 5*10^−6^). (Table 2).

**Table 2.** The total and direct (not mediated via IQ) causal effect estimates of genetic liability to ADHD and ASD on educational attainment.

| **Exposure: genetic liability to ADHD (log-odds). Outcome: educational attainment (SD)** |  |  |  |  |
| --- | --- | --- | --- | --- |
| **Type of effect** | **Beta** | **SE** | **95% CI** | **P-value** |
| Total Effect | -0.103 | 0.023 | -0.15, -0.06 | $5*10^{-6}$ |
| Direct Effect | -0.049 | 0.014 | -0.08, -0.02 | 0.0004 |
| **Exposure: genetic liability to ASD (log-odds). Outcome: educational attainment (SD)** |  |  |  |  |
| **Type of effect** | **Beta** | **SE** | **95% CI** | **P-value** |
| Total Effect | 0.004 | 0.031 | -0.06, 0.07 | 0.9 |
| Direct Effect | 0.028 | 0.013 | 0.002, 0.05 | 0.03 |

The causal effect estimates were directionally consistent across the sensitivity analyses performed. There was limited evidence of horizontal pleiotropy in the analyses, as suggested by the MR Egger intercept (intercept= -0.008; pval= 0.41). Steiger filtering suggested that the direction of the effect was correct for all the ADHD instruments. Table S3 of the supplementary material shows the causal effect estimates, standard errors and p-values derived from the primary and sensitivity analyses.

### Direct effect of genetic liability to ADHD on educational attainment, independent of IQ

The direct causal effect of genetic liability to ADHD on educational attainment (i.e. not via IQ) was approximately 50% smaller than the total causal effect (_MVMR_IVW:-1.6 months per doubling the liability to ADHD; _95%_CI: -2.5 to -0.6; pval= 4*10^−4^) (Table 2). There was evidence of heterogeneity among the causal effect estimates of each instrument included as indicated by the Q statistic (Q= 693; pval= 1*10^−61^).Supplementary table S4 contains the direct causal effect estimates of genetic liability to ADHD and IQ on educational attainment.

### Total causal effect of genetic liability to ASD on educational attainment

The mean F-statistic of the ASD instruments was ≈ 29. There was little evidence of a causal effect of genetic liability to ASD on educational attainment (_MR_IVW: 4 days, per doubling the liability to ASD; _95%_CI: -4.9 months to 5.6 months; pval= 0.9) (Table 2). The confidence intervals across primary and sensitivity analyses were largely overlapping. There was little evidence of directional horizontal pleiotropy (MR Egger intercept= -0.009; pval= 0.42) (Table2). Steiger filtering suggested that the causal effect direction was correct for all the ASD SNPs. Supplementary table S5 contains detailed information on the causal effect estimates, standard errors and p-values across primary and sensitivity analyses.

### Direct effect of genetic liability to ASD on educational attainment, independent of IQ

When allowing for the contribution of IQ, there was some evidence suggesting that a doubling in the genetic liability to ASD has a small positive and direct effect in educational attainment, approximately 29 days (_MVMR_IVW:29 days per doubling the genetic liability to ASD; _95%_CI: 2 to 48; pval= 0.03) (Table 2). There was strong evidence of heterogeneity among the causal effect estimates of each instrument as suggested by the Q statistic (Q= 2380; pval=0).The direct causal effect estimates of genetic liability to ASD and IQ on educational attainment can be found in Supplementary table S6.

### Total causal effect of genetic liability to higher educational attainment on risk of ADHD

The mean F-statistic of the educational attainment instruments was ≈45. There was evidence suggesting that a one SD increase in genetic liability to higher educational attainment (i.e. ≈ 3.8 years of schooling) was associated with approximately 70% lower risk of ADHD (IVW_OR_: 0.30; _95%_CI: 0.26 to 0.36; pval= 6*10^−51^) (Table 3). There was limited evidence of unbalanced horizontal pleiotropy (MR Egger intercept: -0.003; pval= 0.47). Both MR Egger and simex corrected MR Egger estimates, accounting for these pleiotropic effects, were directionally in agreement with the IVW and the confidence intervals across the methods were largely overlapping (Supplementary table S7a). Steiger filtering identified 81 instruments explaining more variation in ADHD than in educational attainment. Removing these instruments attenuated the identified causal effect estimate, which however was still suggestive of a strong causal effect of genetic liability to higher educational attainment on risk of ADHD (Supplementary table S7b).

**Table 3.** The total and direct (not mediated through IQ) causal effect estimates of genetic liability to higher educational attainment on risk of ADHD and ASD diagnosis.

| **Exposure: genetic liability to higher educational attainment (SD). Outcome: ADHD** |  |  |  |  |
| --- | --- | --- | --- | --- |
| Type of Effect | OR | SE | 95% CI | P-value |
| Total Effect | 0.30 | 0.079 | 0.26, 0.36 | $6*10^{-51}$ |
| Direct Effect | 0.33 | 0.126 | 0.26, 0.43 | $6*10^{-17}$ |
| **Exposure: genetic liability to higher educational attainment (SD). Outcome: ASD** |  |  |  |  |
| Type of Effect | OR | SE | 95% CI | P-value |
| Total Effect | 1.51 | 0.082 | 1.29, 1.77 | $5*10^{-7}$ |
| Direct Effect | 1.24 | 0.13 | 0.96, 1.60 | 0.09 |

### Direct effect of genetic liability to higher educational attainment on risk of ADHD, independent of IQ

The estimated causal effect of genetic liability to higher educational attainment on risk of ADHD, independent of IQ, was largely comparable to the total effect (IVW_OR_: 0.33; _95%_CI: 0.26 to 0.43; pval= 6*10^−17^) (Table 3). There was evidence of heterogeneity among the causal effect estimates of each instrument included as indicated by the Q statistic (Q= 843; pval= 2*10^−22^). A direct causal effect was identified even after removing the instruments identified through Steiger filtering (Supplementary table S8). Supplementary table S8, contains the direct causal effect estimates of genetic liability to higher EA and IQ on risk of ADHD.

### Total causal effect of genetic liability to higher educational attainment on risk of ASD

The mean F-statistic of the educational attainment instruments was ≈ 45. There was evidence suggesting that genetic liability to higher educational attainment was associated with increased risk of ASD (IVW_OR_: 1.51 per SD increase; _95%_CI: 1.29 to 1.77; pval= 4*10^−7^) (Table 3). The estimated causal effect was directionally consistent across the sensitivity analyses (Supplementary table S9a) and there was limited evidence to indicate the presence of unbalanced horizontal pleiotropy (MR Egger intercept: -0.007; pval= 0.11). Steiger filtering suggested that 62 SNPs in educational attainment were explaining more variation in ASD and were removed. The exclusion of these SNPs despite attenuating the primary analysis causal effect estimate, was suggestive of a causal effect of genetic liability to higher educational attainment on ASD (Supplementary table S9b).

### Direct effect of genetic liability to higher educational attainment on risk of ASD, independent of IQ

The direct causal effect of genetic liability to higher educational attainment on risk of ASD not mediated through IQ was smaller than the total effect (IVW_OR_: 1.24 per SD increase; _95%_CI: 0.96 to 1.6; pval= 0.09) (Table 3). There was evidence of heterogeneity among the causal effect estimates of each instrument included as indicated by the Q statistic (Q= 910; pval= 2*10^−27^). After removing SNPs identified through Steiger filtering to explain more variation in the outcome, the causal effect estimate attenuated further, providing limited evidence of a direct causal effect of genetic liability to higher educational attainment on risk of ASD (Supplementary table S10). Supplementary table S10 contains the direct causal effect estimates of genetic liability to higher educational attainment and IQ on risk of ASD, as estimated by MVMR analysis.

## DISCUSSION

This is the first study to investigate the bidirectional causal associations between genetic liability to ADHD, ASD and educational attainment and explore the possible role of IQ in the identified causal effects using two-sample MR. Despite the genetic and phenotypic overlap of ADHD and ASD, we found differentiated causal associations of ADHD and ASD genetic liability with educational attainment.

### Bidirectional causal associations between genetic liability to ADHD and educational attainment

We found evidence consistent with a negative causal effect of genetic liability to ADHD on educational attainment, which was only partly attributed to the effects of IQ. This implies that it is ADHD genetic liability itself, not just IQ, that causally influences lower educational attainment. This is in line with a large body of observational evidence suggesting that beyond IQ, ADHD traits and disorder are associated with poor academic outcomes^37,38^. In addition, there is increasing evidence using large cohort and registry data suggesting beneficial effects of ADHD medication on academic performance and outcomes^39,40^. Therefore, our results support and encourage early interventions for ADHD phenotypic expressions in order to improve the academic outcomes of children with ADHD.

We also found that genetic liability to higher educational attainment, over and above IQ, was associated with lower risk of ADHD. This could be explained in the context of educational attainment being associated with several socio-economic status and lifestyle indicators^41,42^. Therefore, our finding supports existing observational evidence suggesting that parental socioeconomic status is associated with risk of ADHD in the offspring^13,43^. This association could be mediated by optimal lifestyle and general health factors during pregnancy, that are known also to be associated with ADHD^44,45^, as well as better prenatal care and access to healthcare services.

Another possible explanation could be that parents with higher educational attainment might place more emphasis on their child’s academic performance, have more access to educational resources and learning stimuli, and cultivate more learning behaviours. In fact, parental resource capital (including income, education and educational material at home) and parental self-efficacy beliefs to help their child, have been found to be important predictors of offspring academic performance^46,47^. Parental emphasis on broader learning behaviours might lead to milder expression, masking or compensation of the ADHD symptomatology in their children, resulting therefore to these children being missed from diagnosis. Academic performance and educational attainment reflect a range of abilities beyond IQ, such as social behaviour^48^, behavioural discipline^49^ and imitation^50^, thus it could be hypothesized that children with genetic liability to higher educational attainment might mask ADHD symptomatology.

### Bidirectional causal associations between genetic liability to ASD and educational attainment

In the case of ASD, we found little evidence suggesting a positive causal effect of genetic liability to ASD on educational attainment. The effect was identified only after the direct, independent of IQ, effects were estimated. In order for this finding to be interpreted, the observational associations of ASD with IQ need to be considered. A large proportion of individuals born with autism present with lower than average IQ and impairments in several areas of neurocognitive functioning^51–53^. The present finding suggests that over and beyond cognitive abilities, phenotypic characteristics of ASD might have small but beneficial effects on educational attainment. Such phenotypic characteristics could include hyper-systemising and attention to detail^54^.

We also identified a positive total causal effect of genetic liability to higher educational attainment on risk of ASD, which MVMR analyses revealed that was attributed, at least partially, to the effects of IQ. This is in line with a recent study utilizing the polygenic transmission disequilibrium test (pTDT) in families of children with ASD, suggesting that parental polygenic risk for higher educational attainment is associated with autism risk in the proband and these probands tend to inherit more alleles associated with higher IQ compared to their neurotypical siblings^55^.

Before reaching conclusions, it is worth considering the extent to which these bidirectional relationships reflect selection bias. Autism diagnosis in the United States has been consistently associated with higher parental socioeconomic status possibly due to better access to health care^56,57^. Although recent evidence from Swedish registry data suggest that the associations between ASD, educational attainment and IQ are unlikely to be influenced by selection bias^14^ future research including samples across countries and socioeconomic strata is necessary.

Overall, in both ADHD and ASD findings, alternative explanations including diagnostic masking and selection bias cannot be rejected since little is currently known on the sociodemographic, socioeconomic and educational factors influencing diagnosis and therefore inclusion in current GWASs.

### Strengths and Limitations

Our study benefitted from using the latest and largest publicly available GWAS data on all the phenotypes of interest. We performed thorough sensitivity analyses to assess the effect of pleiotropic variants used as instruments for each phenotype. We were also able to model the causal effects of each exposure along with IQ so that direct and indirect effects were quantified.

Specific limitations that should be considered. The most important is the use of instruments for ASD below the genome-wide significance threshold (pval< 5*10^−7^). This might have made the ASD analyses prone to weak instrument bias, biasing the estimated causal effect towards the null. However, the estimation of the instruments’ mean F statistic, suggested that this is unlikely.

In addition, there was sample overlap between the educational attainment and IQ GWASs, as both studies included participants from UK Biobank (overlapping participants n= 195,653). This overlap represented approximately 27% of the educational attainment GWAS participants. Overlap between the exposure and outcome GWASs (as in the case of MVMR analyses of ADHD/ASD and IQ on educational attainment) can lead to bias towards the observational estimate^58^. Therefore, for a more accurate estimate of the causal effect of IQ on educational attainment, we orient the readers towards the publication by Davies et al, 2019^59^.

Finally, it is worth considering that ADHD and ASD are highly heterogeneous phenotypes and different phenotypic dimensions have been found to have distinct genetic underpinnings^60,61^. The GWASs utilised in the present study included individuals within the broad range of ADHD and ASD diagnoses, and it is therefore not possible to decipher whether the causal associations identified in the present study are driven by different phenotypic sub-clusters within ADHD and ASD.

### Conclusions and future directions

Despite the genetic and phenotypic overlap of two neurodevelopmental conditions, ADHD and ASD, we found highly differentiated causal associations of ADHD and ASD genetic liability with educational attainment. This emphasizes the necessity for specialised education support services and treatment for affected individuals especially for those with ADHD who seem to be disproportionally affected in terms of their educational outcomes. Further research in necessary in order to elucidate whether the identified causal patterns reflect parentally transmitted effects, diagnostic masking or selection bias and to dissect the broad phenotypes of ADHD, ASD and educational attainment by focusing on investigating causal associations within the several sub-dimensions.

## Funding Acknowledgments

The Medical Research Council (MRC) and the University of Bristol support the MRC Integrative Epidemiology Unit [MC_UU_12013/1, MC_UU_12013/9, MC_UU_00011/1, MC_UU_00011/3, MC_UU_00011/5]. The Economics and Social Research Council (ESRC) support NMD via a Future Research Leaders grant [ES/N000757/1]. CD is funded by the Wellcome Trust [108902/B/15/Z]. BL and LR are supported by the Wellcome Trust (grant ref: 204895/Z/16/Z) awarded to AT, GDS, ES and KT. LDH is supported by a Career Development Award from the UK Medical Research Council [MR/M020894/1] and project entitled ‘social and economic value of health’, which is part of the Health Foundation’s Efficiency Research Programme (grant id: 807293). The Health Foundation is an independent charity committed to bringing about better health and health care for people in the UK. No funding body has influenced data collection, analysis or its interpretation. This publication is the work of the authors, who serve as the guarantors for the contents of this paper. This work was carried out using the computational facilities of the Advanced Computing Research Centre - http://www.bris.ac.uk/acrc/ and the Research Data Storage Facility of the University of Bristol - http://www.bris.ac.uk/acrc/storage/.

